# CRISPR-based surveillance for COVID-19 using genomically-comprehensive machine learning design

**DOI:** 10.1101/2020.02.26.967026

**Authors:** Hayden C. Metsky, Catherine A. Freije, Tinna-Solveig F. Kosoko-Thoroddsen, Pardis C. Sabeti, Cameron Myhrvold

## Abstract

The emergence and outbreak of SARS-CoV-2, the causative agent of COVID-19, has rapidly become a global concern and has highlighted the need for fast, sensitive, and specific tools to surveil circulating viruses. Here we provide assay designs and experimental resources, for use with CRISPR-based nucleic acid detection, that could be valuable for ongoing surveillance. We provide assay designs for detection of 67 viral species and subspecies, including: SARS-CoV-2, phylogenetically-related viruses, and viruses with similar clinical presentation. The designs are outputs of algorithms that we are developing for rapidly designing nucleic acid detection assays that are comprehensive across genomic diversity and predicted to be highly sensitive and specific. Of our design set, we experimentally screened 4 SARS-CoV-2 designs with a CRISPR-Cas13 detection system and then extensively tested the highest-performing SARS-CoV-2 assay. We demonstrate the sensitivity and speed of this assay using synthetic targets with fluorescent and lateral flow detection. Moreover, our provided protocol can be extended for testing the other 66 provided designs. Assay designs are available at https://adapt.sabetilab.org/.

## Introduction

A novel *Severe acute respiratory syndrome-related coronavirus*, SARS-CoV-2 (family: *Coronaviridae*), is the virus behind a severe outbreak originating in China [1]. SARS-CoV-2 surveillance is essential to slowing widespread transmission. In January 2020, we quickly made available a capture enrichment panel [2] using CATCH [3] that is aimed at enhancing sequencing of SARS-CoV-2 and other respiratory viruses. Capture has also been important for ongoing SARS-like coronavirus surveillance [4], and the panel’s inclusion of SARS-like bat and pangolin coronaviruses can aid surveillance efforts.

There are several challenges associated with surveillance during the current SARS-CoV-2 outbreak. First, high case counts overwhelm diagnostic testing capacity, underscoring the need for a rapid pipeline for sample processing [5,6]. Second, SARS-CoV-2 is closely related to other important coronavirus subspecies and species, so detection assays can yield false positives if they are not exquisitely specific to SARS-CoV-2. Third, suspected SARS-CoV-2 patients sometimes have a different respiratory viral infection or have co-infections with SARS-CoV-2 and other respiratory viruses [7]. Therefore, it is important to characterize these other pathogens, for both patient diagnostics and outbreak response.

Here, we help address the challenge of identifying SARS-CoV-2 and the numerous other respiratory viral pathogens by reporting a set of comprehensive design options for 67 species and subspecies for CRISPR-based detection assays. We have not yet experimentally tested most of these designs, instead focusing our efforts so far on extensively testing a point-of-care assay for SARS-CoV-2 using the Cas13-based SHERLOCK technology [6,8,9]. Using this assay, we demonstrate sensitive detection of synthetic SARS-CoV-2 RNA at 10 copies per microliter.

## Results

### Designs for single assay and multiplex panels

We have been developing algorithms and machine learning models for rapidly designing nucleic acid detection assays, linked in a system called ADAPT (manuscript in preparation). The designs satisfy several constraints, including on:

- **Comprehensiveness**: Assays account for a high fraction of known sequence diversity in their species or subspecies (>97% for most assays), and are meant to be effective against variable targets.
- **Predicted sensitivity**: Assays are predicted by our machine learning model to have high detection activity against the full scope of targeted genomic diversity (here, based on *Lwa*Cas13a activity only).
- **Predicted specificity**: Assays have high predicted specificity to their species or subspecies, factoring in the full extent of known strain diversity, allowing them to be grouped into panels that are accurate in differentiating between related taxa.

Comprehensiveness and—to some extent—specificity of the designs can be verified *in silico*.

Using ADAPT we designed 67 assays, satisfying the above constraints, to identify: the SARS-related coronavirus species; SARS-CoV-2; two other subspecies in that species with high similarity to SARS-CoV-2; all other known *Coronaviridae* species, including 4 other species that commonly cause human illness; and other common respiratory viral species and subspecies (**Table 1**). Each assay targets a single species or subspecies and can be used individually (e.g., point-of-care detection); additionally, owing to how they are designed, multiple assays can be grouped together to test for multiple targets and distinguish them with high specificity.

**Table 1.** A summary of the species and subspecies constituting the 67 designs at https://adapt.sabetilab.org/. SARS-CoV-2 is designed to exclude detection of the highly similar RaTG13 sequence, and other similar bat and pangolin SARS-like coronaviruses; the SARS-like subspecies includes most bat and pangolin SARS-like coronaviruses.

| Taxonomic rank | Assay target | Taxonomic rank | Assay target |
| --- | --- | --- | --- |
| Species | *Severe acute respiratory syndrome-related coronavirus* (all known strain diversity) | Species | *Influenza B virus* |
| Subspecies | SARS-CoV-2 | Species | *Human respirovirus 1* (HPIV-1) |
| Subspecies | SARS-CoV-1 | Species | *Human rubulavirus 2* (HPIV-2) |
| Subspecies | SARS-like CoV | Species | *Human respirovirus 3* (HPIV-3) |
| Species | *Human coronavirus 229E* | Species | *Human rubulavirus 4* (HPIV-4) |
| Species | *Human coronavirus NL63* | Species | *Rhinovirus A* |
| Species | *Betacoronavirus 1* (including Human coronavirus OC43) | Species | *Rhinovirus B* |
| Species | *Human coronavirus HKU1* | Species | *Rhinovirus C* |
| Species | *Middle East respiratory syndrome-related coronavirus* | Species | *Enterovirus A* |
| Species | *Influenza A virus* (all subtypes) | Species | *Enterovirus B* |
| Subspecies | H1 (e.g., H1N1 subtype) | Species | *Enterovirus C* |
| Subspecies | H3 (e.g., H3N2 subtype) | Species | *Enterovirus D* |
| Subspecies | N1 (e.g., H1N1 subtype) | Species | *Human orthopneumovirus* (HRSV) |
| Subspecies | N2 (e.g., H3N2 subtype) | Species | *Human metapneumovirus* (HMPV) |
|  |  | Species (39) | All additional species in *Coronaviridae* family |

Sequences for single assays and multiplex panels are available at **https://adapt.sabetilab.org/**.

### SARS-CoV-2 SHERLOCK assay testing

We initially screened a set of 4 designs for SHERLOCK [6,8,9] assays, output by ADAPT to detect SARS-CoV-2. We identified an assay, which was the best-performing and also our highest ranked design *a priori*. We extensively tested this assay using a synthetic RNA target and determined the limit of detection to be 10 copies/µl using both fluorescent and lateral flow detection (**Figure 1**). This assay performs well in comparison to the recently disclosed DETECTR [10] assay (sensitivity: 70–300 cp/µl) [11] and SHERLOCK assay (10–100 cp/µl) [12] for SARS-CoV-2. A protocol for performing this assay is provided in the Methods section and can be used for testing any of the other designs we have provided.

**Figure 1.**
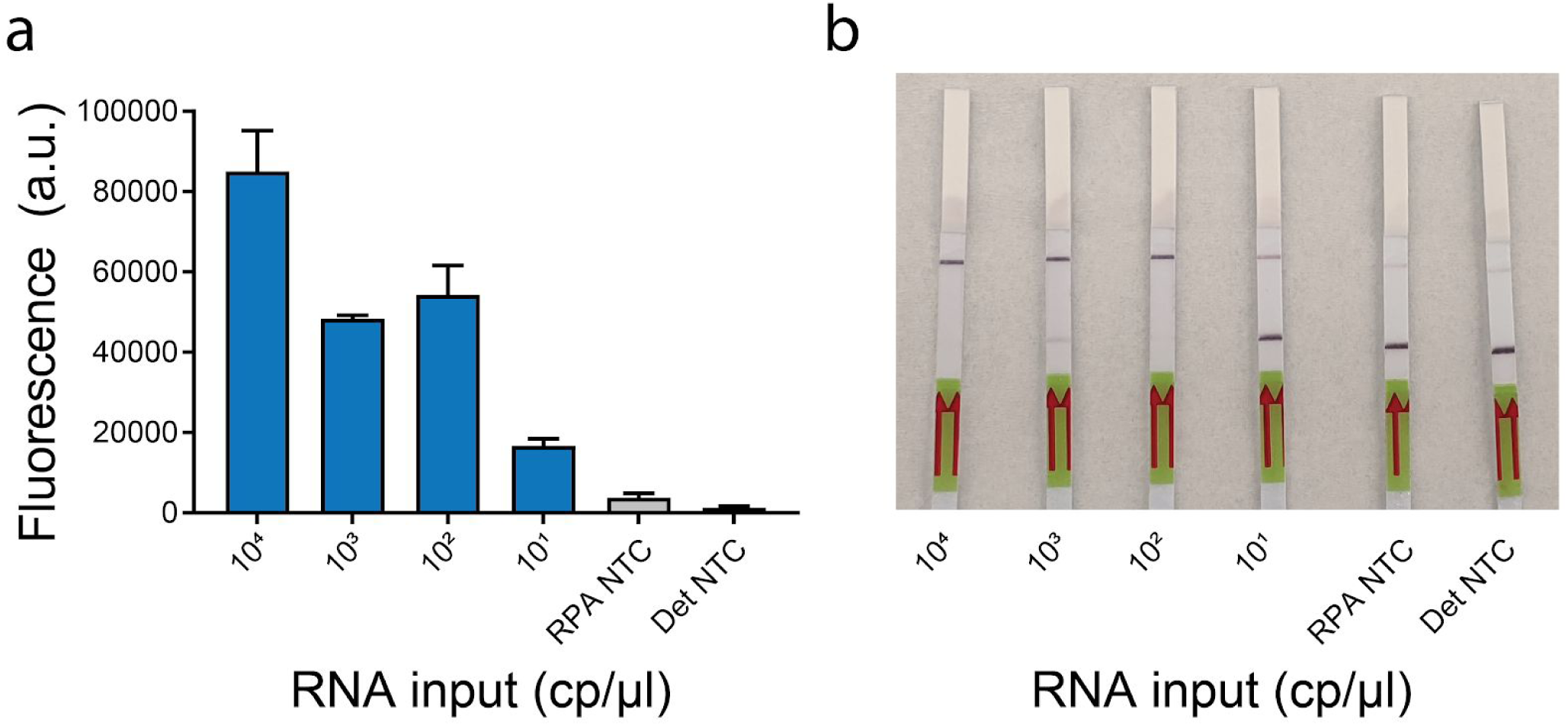
Testing an assay for SARS-CoV-2 using synthetic RNA targets. We show data from both fluorescent (a) and lateral flow detection (b). Target concentrations (in cp/μl) are indicated. RPA NTC: water input into RPA; Det NTC: water input into Cas13 detection reaction. In (a), error bars indicate one standard deviation based on *n*=3 technical replicates.

## Discussion

Ongoing SARS-CoV-2 sequencing is key to developing and monitoring diagnostics and similar surveillance tools. In the case of the SARS-CoV-2 outbreak, genomes have been generated and shared at a remarkable pace, and we thank those who have contributed their data through GISAID [13]. We and others, relying on this data [14], have shown that it is possible to rapidly design CRISPR-based tools for detection and surveillance during an outbreak.

Among other goals for this work, we plan to evaluate: (1) sensitivity of the SARS-CoV-2 assay against clinical isolates and patient samples—including sputum, throat, and nasal swabs—some of which may be challenging sample types to test; (2) specificity at both the species and subspecies levels against highly related viruses. For the latter, we intend to use a mixture of synthetic targets reflecting different viral sequences, and patient samples or viral seedstocks when available. We hope that the comprehensiveness and high predicted sensitivity and specificity of our designs will enable many groups to proceed rapidly and successfully from assay testing through deployment.

## Acknowledgements

We thank Brittany Petros, Christopher Tomkins-Tinch, Molly Kemball, and Megan Vodzak for their contributions and help, and other members of the Sabeti lab for helpful conversations related to this work. Funding was provided by HHMI and DARPA D18AC00006.

We also wish to thank the following labs that have generously made available SARS-CoV-2 samples and genome sequences via GISAID:

- Amalea Dulcene Nicolasora Research Institute for Tropical Medicine, Molecular Biology Laboratory
- Arizona Department of Health Services
- Bamrasnaradura Hospital
- Beijing Genomics Institute (BGI)
- Beijing Institute of Microbiology and Epidemiology
- BGI & Institute of Microbiology, Chinese Academy of Sciences & Shandong First Medical University & Shandong Academy of Medical Sciences & General Hospital of Central Theater Command of People’s Liberation Army of China
- California Department of Public Health
- Centers for Disease Control, R.O.C. (Taiwan)
- Centre for Infectious Diseases and Microbiology Laboratory Services
- Charité Universitätsmedizin Berlin, Institute of Virology
- Chongqing Municipal Center for Disease Control and Prevention
- CNR Virus des Infections Respiratoires - France SUD
- Collaboration between the University of Melbourne at The Peter Doherty Institute for Infection and Immunity, and the Victorian Infectious Disease Reference Laboratory
- Department of Disease Control, Ministry of Public Health, Thailand
- Department of Infectious and Tropical Diseases, Bichat Claude Bernard Hospital, Paris
- Department of Laboratory Medicine, Lin-Kou Chang Gung Memorial Hospital, Taoyuan, Taiwan.
- Department of Laboratory Medicine, National Taiwan University Hospital
- Department of Medical Sciences, Ministry of Public Health, Thailand
- Department of Microbiology, Guangdong Provincial Center for Diseases Control and Prevention
- Department of Microbiology, Zhejiang Provincial Center for Disease Control and Prevention
- Department of Virology III, National Institute of Infectious Diseases
- Department of Virology, University of Helsinki and Helsinki University Hospital, Helsinki, Finland
- Dept. of Pathology, National Institute of Infectious Diseases
- Division of Viral Diseases, Centers for Disease Control and Prevention
- Fujian Center for Disease Control and Prevention
- General Hospital of Central Theater Command of People’s Liberation Army of China
- Guangdong Provincial Center for Diseases Control and Prevention; Guangdong Provincial Public Health
- Hangzhou Center for Disease Control and Prevention
- Hong Kong Department of Health
- Hubei Provincial Center for Disease Control and Prevention
- Illinois Department of Public Health Chicago Laboratory
- INMI Lazzaro Spallanzani IRCCS
- Institut für Mikrobiologie der Bundeswehr, Munich
- Institute of Pathogen Biology, Chinese Academy of Medical Sciences & Peking Union Medical College
- Institute of Viral Disease Control and Prevention, China CDC
- Jiangsu Provincial Center for Disease Control & Prevention
- Joanna Ina Manalo Research Institute for Tropical Medicine, Molecular Biology Laboratory
- Korea Centers for Disease Control & Prevention (KCDC) Center for Laboratory Control of Infectious Diseases Division of Viral Diseases
- KU Leuven, Clinical and Epidemiological Virology
- Laboratoire Virpath, CIRI U111, UCBL1, INSERM, CNRS, ENS Lyon
- Laboratory of Virology, INMI Lazzaro Spallanzani IRCCS
- Lapland Central Hospital
- Li Ka Shing Faculty of Medicine, The University of Hong Kong
- Massachusetts Department of Public Health
- Microbial Genomics Core Lab, National Taiwan University Centers of Genomic and Precision Medicine
- Monash Medical Centre
- National Centre for Infectious Diseases
- National Influenza Centre, National Public Health Laboratory, Kathmandu, Nepal
- National Institute for Communicable Disease Control and Prevention (ICDC) Chinese Center for Disease Control and Prevention (China CDC)
- National Institute for Viral Disease Control & Prevention, China CDC
- National Public Health Laboratory, National Centre for Infectious Diseases
- National Reference Center for Viruses of Respiratory Infections, Institut Pasteur, Paris
- NHC Key laboratory of Enteric Pathogenic Microbiology, Institute of Pathogenic Microbiology
- NSW Health Pathology - Institute of Clinical Pathology and Medical Research; Westmead Hospital; University of Sydney
- Pamela Youde Nethersole Eastern Hospital
- Pathogen Discovery, Respiratory Viruses Branch, Division of Viral Diseases, Centers for Disease Control and Prevention
- Pathogen Genomics Center, National Institute of Infectious Diseases
- Pathology Queensland
- Prince of Wales Hospital (PWH)
- Princess Margaret Hospital (PMH)
- Programme in Emerging Infectious Diseases, Duke-NUS Medical School
- Providence Regional Medical Center
- Public Health Virology Laboratory
- Queen Elizabeth Hospital (QEH)
- Queen Mary Hospital (QMH)
- Respiratory Virus Unit, Microbiology Services Colindale, Public Health England
- Serology, Virology and OTDS Laboratories (SAViD), NSW Health Pathology Randwick
- Shenzhen Key Laboratory of Pathogen and Immunity, National Clinical Research Center for Infectious Disease, Shenzhen Third People’s Hospital
- Singapore General Hospital, Molecular Laboratory, Division of Pathology
- Sorbonne Université, Inserm et Assistance Publique-Hôpitaux de Paris (Pitié Salpêtrière)
- South China Agricultural University
- State Key Laboratory of Virology, Wuhan University
- Taiwan Centers for Disease Control
- Texas Department of State Health Services
- Thai Red Cross Emerging Infectious Diseases - Health Science Centre
- The University of Hong Kong - Shenzhen Hospital
- Tuen Mun Hospital (TMH)
- Unit for Laboratory Development and Technology Transfer, Public Health Agency of Sweden
- United Christian Hospital
- Virology Laboratory National Institute for Infectious Diseases ‘Lazzaro Spallanzani’ IRCCS
- Virology Laboratory, INMI L. Spallanzani
- Virology Unit, Institut Pasteur du Cambodge
- Washington State Department of Health
- Wisconsin Department of Health Services
- Wuhan Fourth Hospital
- Wuhan Institute of Virology, Chinese Academy of Sciences
- Wuhan Jinyintan Hospital
- Yongchuan District Center for Disease Control and Prevention
- Zhejiang Provincial Center for Disease Control and Prevention
- Zhongxian Center for Disease Control and Prevention

## Conflicts of interest

H.C.M., C.A.F., P.C.S., and C.M. are inventors on patent filings related to this work. P.C.S. is a co-founder of, shareholder in, and advisor to Sherlock Biosciences, Inc, as well as a Board member of and shareholder in Danaher Corporation.

## Methods

### SHERLOCK protocol for SARS-CoV-2

#### Isothermal amplification (RT-RPA)

##### List of equipment and materials

- Heat block, water bath, or thermocycler, prewarmed to 41 °C
- RevertAid Reverse Transcriptase (Thermo)
- RNase inhibitor (NEB Murine)
- Primer mix @ 5 µM of each primer (see Supplementary Table 1 for primer sequences)
- Synthetic DNA or RNA target (see Supplementary Table 1 for sequences), or extracted RNA from a viral seedstock or patient sample
- Rehydration buffer, lyophilized RPA pellets, MgAc @ 280 mM from RPA kit (TwistAmp Basic kit, TwistDx)
- Nuclease-free water

| Reagent | Initial concentration | Amount to add for N RPA pellets |
| --- | --- | --- |
| Rehydration buffer | N/A | $29.5 \times N \mu\text{l}$ |
| RPA primer mix | 5 µM of each primer | $4.8 \times N \mu\text{l}$ |
| Nuclease-free water | N/A | $2.1 \times N \mu\text{l}$ |
| RNase inhibitors | 40 U/µl | $5 \times N \mu\text{l}$ |
| RevertAid RT | 200 U/µl | $N \mu\text{l}$ |
| Magnesium Acetate (MgAc) | 280 mM | $3.04 \times N \mu\text{l}$ |

##### Step by step protocol

1. Determine the number of pellets needed based on the size of the experiment (2 samples per pellet, if doing 20 µl RPA reactions).
2. Make a master mix for N pellets, consisting of Rehydration buffer, RPA primer mix, water, RNase inhibitors, and RevertAid RT. Do not include MgAc at this step.
3. Resuspend each RPA pellet using the total volume for 1 RPA pellet (∼30 µl) of master mix.
4. Add the MgAc to the master mix. Keep the master mix on ice.
5. Aliquot the master mix (containing MgAc) into wells of a 96-well plate or strip tubes pre-chilled on ice. For 20 µl reactions, aliquot 18 µl of master mix.
6. Add sample or a control target to each aliquot of master mix (2 µl, if using 20 µl reactions), mix thoroughly, and incubate at 41 °C for 20 minutes.

#### Cas13 detection

##### List of equipment and materials

- Heat block, water bath, or thermocycler, prewarmed to 37 °C
- **For visual readout:** camera or cell phone with camera; **for fluorescent readout:** qPCR machine / plate reader capable of detecting FAM
- **For fluorescent readout:** optical 96-well plate, optical strip-tubes, or black 96-well plate with clear bottom
- 10× Cleavage buffer (1× CB is 40 mM Tris pH 7.5, 1 mM DTT)
- RNase inhibitors (NEB Murine)
- Cleavage reporter, **for visual readout:** IDT for lateral flow (sequences in Supplementary Table 1); **for fluorescent readout:** RNase Alert v2 (Thermo)
- *Lwa*Cas13 protein @ 0.5 mg/ml, in 16 ul aliquots, diluted in 1× Storage buffer (50 mM Tris pH 7.5, 600 mM NaCl, 5% glycerol, 2 mM DTT) [15]
- Cas13 crRNA @ 2 µM (see Supplementary Table 1 for sequences)
- T7 RNA polymerase (Lucigen NxGen)
- rNTP mix @ 25 mM each (NEB)
- MgCl_2_ @ 100 mM
- Nuclease-free water

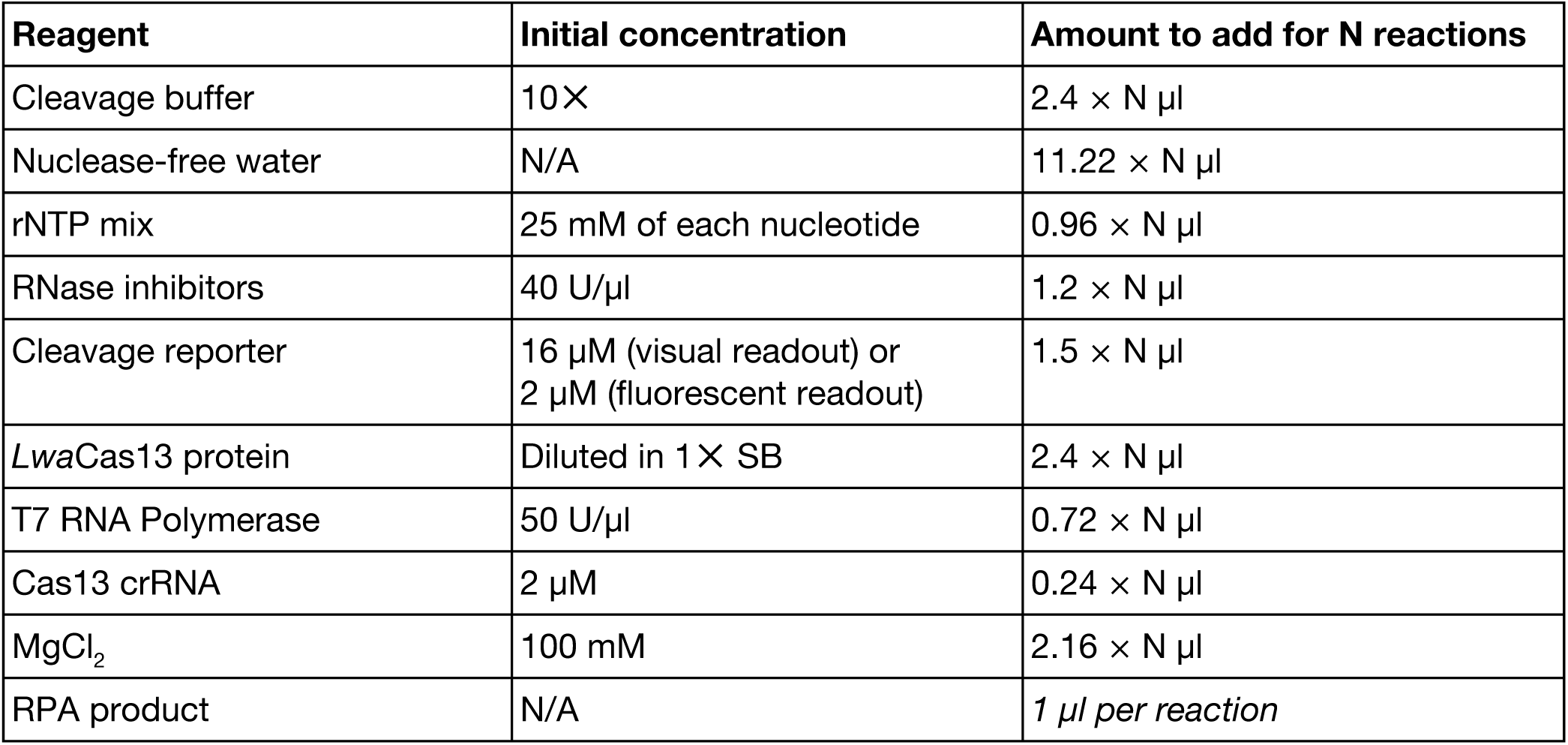

##### Step by step protocol

1. Dilute *Lwa*Cas13a by adding 110.5 µl of 1× Storage buffer to a 16 µl aliquot of Cas13 protein (prior to dilution @ 0.5 mg/ml).
2. Prepare a master mix based on the table above, adding the components in the order they are listed in the table. Do not add RPA product at this step.
3. Aliquot 19 µl of master mix into wells of a 96-well plate or strip tubes pre-chilled on ice. If using a fluorescent readout and depending on the instrument used, an optical PCR or black plate with a clear base can be used.
4. Add 1 µL of RPA product to each master mix aliquot. Seal the plate and incubate at 37 °C for 30 minutes to 3 hours. **For fluorescent readout**, measure fluorescence every 5 minutes. **For visual readout**, see additional details below.

#### Visual readout

##### List of equipment and materials

- HybriDetect 1 lateral flow strips (Milenia)
- Hybridetect Assay Buffer (Milenia)

##### Step by step protocol

1. After incubation at 37 °C for 30 minutes to 3 hours, add 80 µl of Hybridetect Assay Buffer to the total volume of each detection reaction.
2. Add a lateral flow strip to each well and incubate for 2-5 minutes at room temperature.
  A. Take care to avoid contamination of the strips, by using tweezers to remove individual strips and place in buffer.
3. Remove strips, place on flat, well-lit surface, and analyze or acquire images of the strips.

## Supplementary Information

**Supplementary Table 1.**
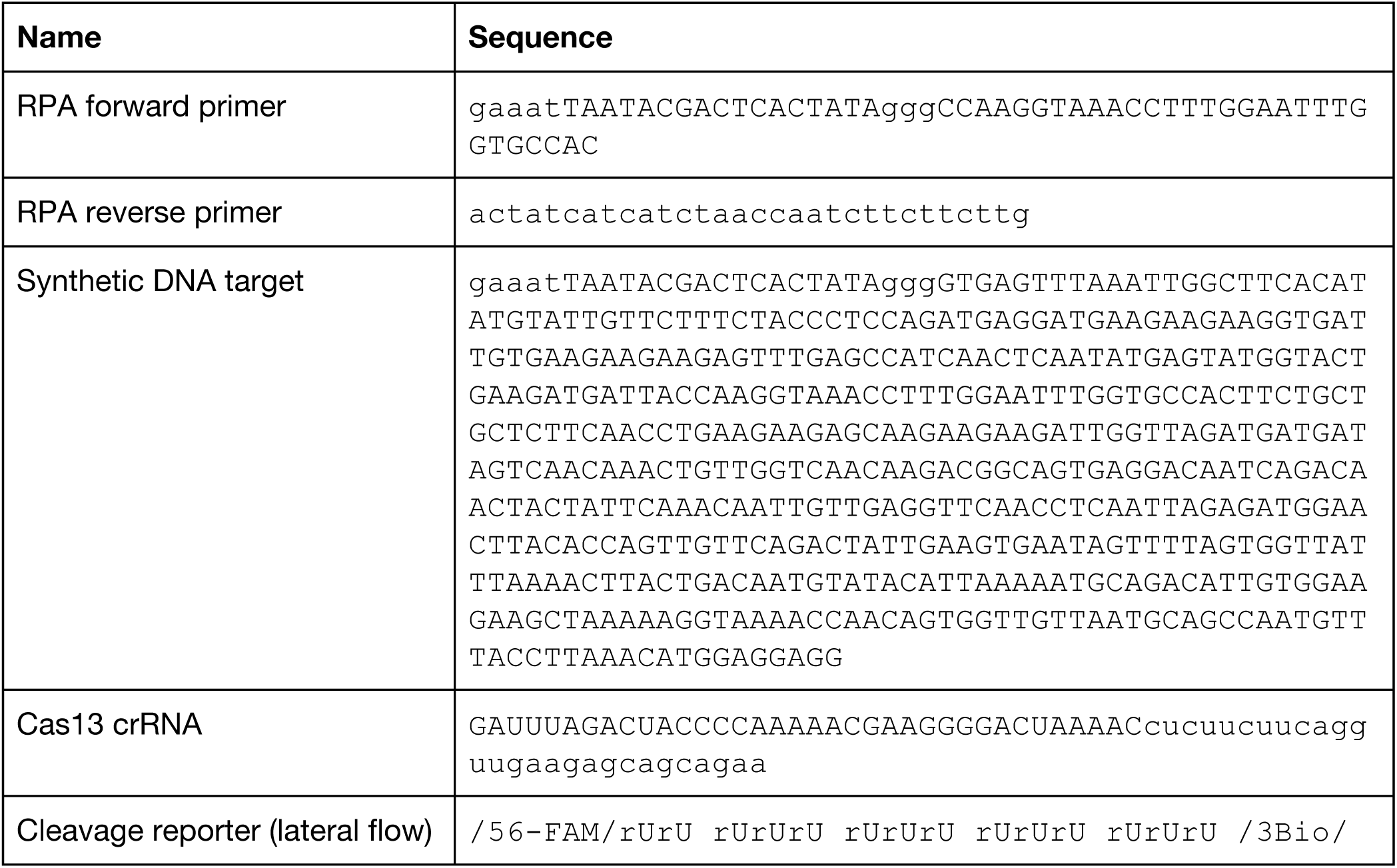
Primer, target, and crRNA sequences for the SARS-CoV-2 assay.

## Notes

#### Summary of Updates

Amended title; revised language around diagnostics to more accurately reflect current state; fixed issue with links.

https://adapt.sabetilab.org

## References

1. Zhou P, Yang X-L, Wang X-G, Hu B, Zhang L, Zhang W, et al. A pneumonia outbreak associated with a new coronavirus of probable bat origin. Nature. 2020. doi:10.1038/s41586-020-2012-7

2. V-Respiratory probe set (2020-01). Available: https://github.com/broadinstitute/catch/tree/master/probe-designs

3. Metsky HC, Siddle KJ, Gladden-Young A, Qu J, Yang DK, Brehio P, et al. Capturing sequence diversity in metagenomes with comprehensive and scalable probe design. Nat Biotechnol. 2019;37: 160–168. doi:10.1038/s41587-018-0006-x

4. Li B, Si H-R, Zhu Y, Yang X-L, Anderson DE, Shi Z-L, et al. Discovery of Bat Coronaviruses through Surveillance and Probe Capture-Based Next-Generation Sequencing. mSphere. 2020;5. doi:10.1128/mSphere.00807-19

5. Wee S-L. As Deaths Mount, China Tries to Speed Up Coronavirus Testing. The New York Times. 9 Feb 2020. Available: https://www.nytimes.com/2020/02/09/world/asia/china-coronavirus-tests.html. Accessed 24 Feb 2020.

6. Myhrvold C, Freije CA, Gootenberg JS, Abudayyeh OO, Metsky HC, Durbin AF, et al. Field-deployable viral diagnostics using CRISPR-Cas13. Science. 2018;360: 444–448. doi:10.1126/science.aas8836

7. Wang M, Wu Q, Xu W, Qiao B, Wang J, Zheng H, et al. Clinical diagnosis of 8274 samples with 2019-novel coronavirus in Wuhan. Infectious Diseases (except HIV/AIDS). medRxiv; 2020. doi:10.1101/2020.02.12.20022327

8. Gootenberg JS, Abudayyeh OO, Lee JW, Essletzbichler P, Dy AJ, Joung J, et al. Nucleic acid detection with CRISPR-Cas13a/C2c2. Science. 2017;356: 438–442. doi:10.1126/science.aam9321

9. Gootenberg JS, Abudayyeh OO, Kellner MJ, Joung J, Collins JJ, Zhang F. Multiplexed and portable nucleic acid detection platform with Cas13, Cas12a, and Csm6. Science. 2018;360: 439–444. doi:10.1126/science.aaq0179

10. Chen JS, Ma E, Harrington LB, Da Costa M, Tian X, Palefsky JM, et al. CRISPR-Cas12a target binding unleashes indiscriminate single-stranded DNase activity. Science. 2018;360: 436–439. doi:10.1126/science.aar6245

11. James P. Broughton, Wayne Deng, Clare L. Fasching, Jasmeet Singh, Charles Y. Chiu, Janice S. Chen. A protocol for rapid detection of the 2019 novel coronavirus SARS-CoV-2 using CRISPR diagnostics: SARS-CoV-2 DETECTR. 2020. Available: https://mammoth.bio/wp-content/uploads/2020/02/A-protocol-for-rapid-detection-of-the-2019-novel-coronavirus-SARS-CoV-2-using-CRISPR-diagnostics-SARS-CoV-2-DETECTR.pdf

12. Feng Zhang, Omar O. Abudayyeh, Jonathan S. Gootenberg. A protocol for detection of COVID-19 using CRISPR diagnostics. Available: https://www.broadinstitute.org/files/publications/special/COVID-19%20detection%20(updated).pdf

13. Shu Y, McCauley J. GISAID: Global initiative on sharing all influenza data - from vision to reality. Euro Surveill. 2017;22. doi:10.2807/1560-7917.ES.2017.22.13.30494

14. Hou T, Zeng W, Yang M, Chen W, Ren L, Ai J, et al. Development and Evaluation of A CRISPR-based Diagnostic For 2019-novel Coronavirus. Infectious Diseases (except HIV/AIDS). medRxiv; 2020. doi:10.1101/2020.02.22.20025460

15. Kellner MJ, Koob JG, Gootenberg JS, Abudayyeh OO, Zhang F. SHERLOCK: nucleic acid detection with CRISPR nucleases. Nat Protoc. 2019;14: 2986–3012. doi:10.1038/s41596-019-0210-2

